# A novel simultaneous RNA extraction workflow from mouse Retinal Pigmented Epithelial (RPE) and retina

**DOI:** 10.64898/2026.11.25.769171

**Authors:** Shomoita Sayed, Philip Low, Madduri Srinivasarao

**Author notes:** Funding: Funding from the Low Institute of Therapeutics (LIFT) and Purdue University. Manuscript Word Count: 2297 (excluding the structured abstract). Commercial relationships disclosure: N (No commercial financial relationships) Special Comments on authors: Dr. Philip Low passed away before submission.

## Abstract

**Purpose:** In Posterior Segment Eye Diseases (PSEDs), the homeostatic relation between retinal pigment epithelium (RPE) and retina is disrupted. A better understanding of the transcriptomic changes occurring in different ocular cell types can help to elucidate disease mechanisms. Currently, there is no complete detailed workflow describing the process of simultaneous RNA isolation from RPE and retina separately. In this paper, our aim was to develop an easy, reproducible protocol for simultaneous isolation of RNA from both murine RPE and retina.

**Methods:** We developed an efficient and straightforward workflow for isolating RNA from both murine RPE and retina simultaneously. We described the method in detail spanning from mouse euthanasia to downstream RNA analysis. We compared the efficiency of this isolation method between different mice strains (C57BL/6 vs BALB/c).

**Results:** Our method provided high yield of RNA from both RPE and retina across different strains. We observed better quality of RPE RNA in terms of A260/130 ratios in C57BL/6 mice. The retinal RNA quality and yield were comparable across both strains. Beta-catenin immunofluorescent staining also validated our success of RPE removal in the choroidal flat-mounts in both strains.

**Conclusions:** We described an improved workflow for obtaining RNA simultaneously from both RPE and retina. We compared the efficiency of our process via multitude of downstream analyses across different mice strains.

**Translational Relevance:** This method will be beneficial to understand the RPE or retinal tissue specific mRNA transcript changes in preclinical mice models of PSEDs.

## Introduction

Posterior Segment Eye diseases (PSED)s such as Age-related Macular Degeneration (AMD) and Diabetic retinopathy affected approximately 299 million people globally in 2020 [1-2]. The number of people suffering from PSEDs is estimated to be over 440 million by 2040 [1-2]. An in-depth understanding of the disease manifestation across different ocular cell populations can facilitate the development of novel therapies for the PSEDs. In PSEDs, the homeostatic functions of Retinal Pigmented Epithelial (RPE) cells and retina are dysregulated [3-7]. The posterior segment of the eye majorly comprises of RPE and retina [8-9]. The interconnected relation between RPE and retinal cells plays an important role in the visual cycle and healthy ocular homeostasis. The RPE, a single monolayer of cells located between the retina and choroidal vasculature, creates the selective outer blood-retinal-barrier (BRB) and supports the retinal homeostasis [8]. The neurosensory retina transduces the light signal into electrochemical signal for visual cycle [9]. Understanding the cell specific transcriptomic changes can provide important information about the overall disease pathogenesis. Mice models are the standard disease models to understand complex disease pathologies [10-11]. But due to the small eye size, fragility and tissue structure of murine eye, the simultaneous isolation of RPE and retinal RNA can pose difficulties. Preclinical studies often rely on complicated methods such as Laser capture microdissection (LCM) to isolate pure cell populations to understand the cell specific genetic changes [12-13]. But LCM is an expensive and time-consuming process. Mechanically or enzymatically digested workflow for RPE RNA isolation has been described from rat eyes but not mouse eyes [14-15].

Here we describe a time-efficient way of simultaneous RNA isolation from both murine RPE and the retina for downstream RNA analysis.

## 2. Material and Methods

### 2.1: Animals

Adult C57BL/6 and BALB/c mice used for these experiments were purchased from Charles River and Jackson Laboratory, respectively. The mice were kept at animal housing facility in Purdue University under normal lighting conditions with 12 h on, 12 h off cycles. All animal procedures were performed in accordance with the Association for Research in Vision and Ophthalmology (ARVO) Statement for the Use of Animals in Ophthalmic and Vision research and were approved by the institutional animal care and use committee (IACUC) of Purdue University.

### 2.2: Simultaneous RPE and Retinal RNA isolation

Enucleation & dissection: The mice were euthanized using carbon dioxide asphyxiation followed by secondary cervical dislocation. For enucleation, gentle pressure was applied on the skin surrounding the eye, which caused the outward protrusion of the eyeball. Using forceps, the eyeball was carefully removed from the ocular cavity. Then the collected eye was readily submerged into the RNA-protect solution. Preservation of the eyeball in the solution enabled isolation of the RNA in the future. 200-300 µl of the preserving solution is sufficient for submerging one murine eyeball. An eyeball preserved in RNA-protect solution can be dissected instantly or stored for RNA isolation at a later time-point. At the time of dissection, the eyeball was transferred to the cold PBS in a round petri dish. The extraocular tissue and muscles were excised using spring scissors. The optic nerve was kept intact to enable easy maneuvering of the eyeball. Using forceps the cornea was pinched forward to create a small incision. Subsequently, the cornea was carefully excised, and the iris, ciliary bodies, and lens were removed. After removal of lens, a small slit was created using the spring scissor in the posterior part of the eye. To achieve clean separation of the choroid-RPE and retina, the forceps were gently inserted into the slit and moved circularly along the half-circular posterior segment to detach the retina from the Choroid-RPE. The retina was removed from the choroid using forceps.

Simultaneous RNA isolation from RPE and Retina: Once the retina and choroid-RPE were separated, the process of simultaneous RNA isolation was started. The separated sclera-choroid-RPE and retina were transferred to a 1.5 mL centrifuge tubes containing 200 & 400 µl of RLT buffer with beta-Mercaptoethanol (B-ME), respectively. (Note: Due to the hazardous nature of B-ME, the user should wear proper protective face masks and conduct steps involving RLT buffer inside a fume hood). Carrier RNA (provided by the Qiagen Kit) was used to increase the yield of RNA.

Addition of carrier RNA was particularly important for maximizing RNA yield from the RPE due to the small size of the RPE cell population. Accordingly, 20 ng of carrier RNA was added to each tube. The tubes were vigorously shaken for 3-5 seconds for a period of 7-10 minutes. The shaking ensured the dispersion of RPE cells in the RLT buffer. The successful RPE cell digestion was indicated by the color change of the RLT solution. A sterile pestle attached to a hand-held tissue homogenizer was used for the complete digestion of retina. With the gentle up-and-down strokes of the pestle, the tissue was homogenized. Excessive pressure was avoided to prevent foam formation, which could result in tissue loss. An equal volume of 70% ethanol was added to each tube. The tubes were flicked and gently shaken to ensure proper mixing of the ethanol with the extraction buffer. The solution was then transferred to an RNeasy MinElute spin column containing collection tubes. The tubes were centrifuged for 1-3 minutes at 15000 g. Followed by the disposal of the RLT flow-through, 350 µl Buffer RW1 was carefully added to the spin columns and centrifuged at the same speed. After the removal of RWI flow-through, 80 µl of DNase buffer stock (DNase: RDD buffer= 1:8) was added carefully to the spin columns. The tubes were then placed at room temperature for 15 min. An additional 350 µl Buffer RW1 was added to the spin columns before a second round of centrifugation. After the second RW1 buffer wash, the spin columns were transferred into new collection tubes to minimize the risk of contamination. 500 µl Buffer RPE was added to the spin columns. The tubes were then centrifuged at the same settings. After discarding the RPE buffer flow-through, 500 µl of 80% ethanol was added to the spin columns and centrifuged at the same settings. A second optional ethanol wash was conducted. To ensure the complete removal of residual ethanol from the spin column membranes, the tubes were centrifuged for 5 minutes with lids kept open. In the final step, the spin columns were carefully placed in RNase free tubes. To elute the RNA, 17-18 µl of RNase-free water was directly added to the center of the spin column membrane, followed by an incubation at the room temperature for at least 5 minutes. After a final centrifuge for 2 mins at 15000 g, the spin columns were removed and the RNA containing tubes were immediately transferred to ice to avoid RNA degradation. Several of the steps of the workflow are shown in Figure-1 to provide the readers with a better understanding. The reagents used in the extraction workflow and the downstream analyses are also provided in table-1.

**Table 1:** Reagents used during dissection & RNA isolation.

| **Material** | **Brand** | **Catalog** |
|---|---|---|
| Forceps | Fine Science Tools | FST 11251-20 |
| Spring scissor | Fine Science Tools | FST91500-09 |
| RNase free Centrifuge tube | INVITROGEN | AM12400 |
| Petri dish | Fisher Scientific | FB087571 |
| PBS | Corning | 21-040-cv |
| RNA-protect solution | Qiagen | 76104 |
| RNeasy Micro Kit | Qiagen | 74004 |
| DNase enzyme | Qiagen | 79254 |
| Hand-held tissue homogenizer | Fisher Scientific | 12-141-361 |
| RNase-free disposable pellet pestles | Fisher Scientific | 12-141-364 |
| Beta-Mercapto-ethanol | Sigma Aldrich | 444203-250ML |

**Figure 1:**
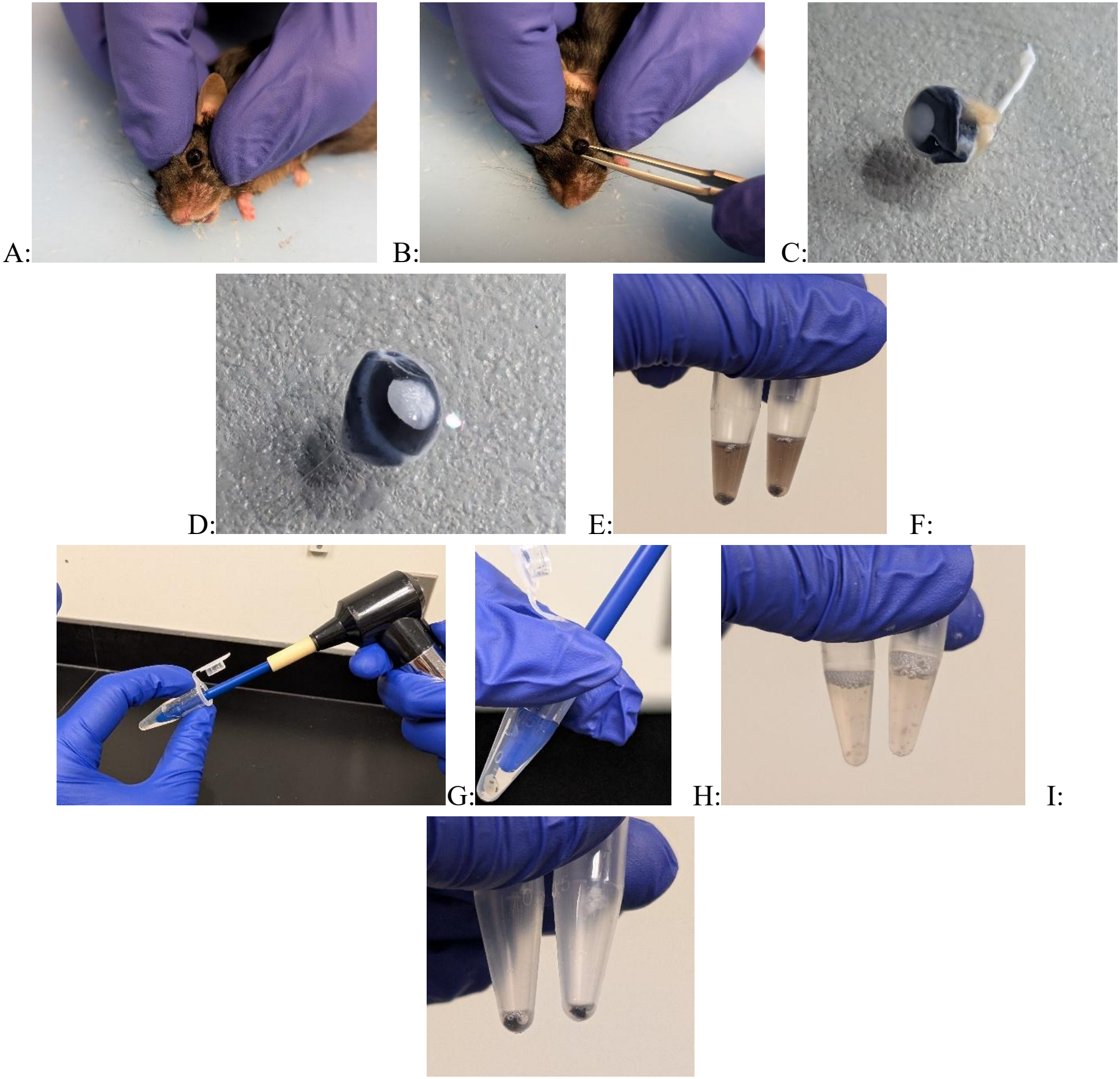
(A-I): Workflow for simultaneous RNA isolation from murine RPE and retina. A: the eyeball is protruding due to the pressure applied around the surrounding skins, B: The fine tips of the forceps are placed beneath the eyeball, C: Eyeball with the optic nerve and extraocular tissue is preserved in the RNA-protect solution, D: Cleaned eyeball, E: The color of the Lysis buffer changes after RPE cells are released, F: Hand-held homogenizer is used for the retinal homogenization, G: Magnified image of the retina, H: Lysis buffer gets cloudy after retinal tissue homogenization, I: The choroid tissue remains at the bottom of centrifuge tubes after removal of RPE containing lysis buffer.

### 2.3. Assessment of ocular RNA yield

To assess the yield, 1.2 µl of isolated RNA was used. Using Nanodrop spectrophotometer, the RNA yield, A260/280 and A260/230 ratio were quantified for each sample. The RNA was then stored in -80 °C. Welch’s T test was conducted using GraphPad Prism 11.0.2 to compare the yield and purity ratios between the two strains.

### 2.4. Immunostaining of whole flat-mount choroid

To ensure the success of the isolation method, immunofluorescent staining on choroids was conducted. The choroid tissue was fixed using 4% Paraformaldehyde (PFA) for 1 hour followed by permeabilization with 0.1% (10 minutes) and 1.5 % PBST (30 minutes), respectively. Then the choroid tissue was blocked with blocking buffer (10% goat serum in sterile PBS) overnight at 4 °C. Followed by the blocking step, the tissue was incubated with beta-catenin [Genetex: GTX-1435] for 24 hours using the same blocking buffer. After primary antibody incubation, the tissue was permeabilized again using 0.1% PBST for 30 minutes followed by incubation with secondary antibody [Invitrogen: A-21244] overnight. In the final step, the tissue was washed in PBS for 5 minutes and then mounted on charged slides with Fluor mount G mounting medium. The images were taken at 20X magnification using the Zeiss LSM 900 microscope. In table-2, the reagent information is provided.

**Table 2:** Reagents used for downstream validation experiments.

| **Reagents** | **Brand** | **Catalog** |
|---|---|---|
| Triton-X-100 | Thermo Scientific Chemicals | AC215680025 |
| Goat serum | Invitrogen |  |
| Beta-catenin | Genetex | GTX01435 |
| Goat anti-Rabbit IgG (H+L) Secondary Antibody, Alexa Fluor™ 647 | Thermofisher Scientific | A-21244 |
| Fluoromount-G mounting medium | Invitrogen | 00-4958-02 |
| Microscopic slides | Polyscience Inc. | 24216 |
| Coverslips | Polyscience Inc. | 24071-1 |
| Disposable transfer pipette | Fisher Scientific | 16001-188 |
| 12 well plates | Fisher Scientific | FB012928 |
| PCR reagents |  |  |
| Universal SYBR Green One-step kit | Bio-RAD | 1725151 |
| Hard-shell PCR plates | Bio-rad | HSP9601 |
| RNase free Centrifuge tube | Invitrogen | AM12400 |

### 2.5 Biomarker analysis using RT-PCR

To assess the quality of our isolation techniques, we conducted qPCR. In RPE samples, we compared RPE65 (RPE biomarker) and Collagen-6 (choroid capillaries biomarker). To check the purity of retinal tissue, Rhodopsin (Retinal Marker) and RPE65 were assessed. The primers were designed by IDT-DNA following the already described primer sequences [16-17]. Using Bio-Rad One-step Universal SYBR green kit, 20 nanogram of template RNA was used in 10 µl PCR volume per well. The PCR reaction was conducted as per the kit manual. Each sample wasrun at triplicates. The average Cq values were normalized to housekeeping gene, GAPDH. Then the normalized values were assessed against tissue specific respective biomarkers to generate the relative mRNA fold changes. Welch’s T test was conducted using GraphPad Prism 11.0.2.

## 3. Results

### 3.1. RNA yield and purity

In terms of RPE derived RNA, C57BL/6 and BALB/c showed similar yields (Figure-2). The RNA purity as indicated by the 260/280 ratio was also comparable between two strains as shown in Figure-2. But the 260/230 ratio was significantly better in the C57BL/6. The range of the 260/230 ratio indicated presence of chemical contaminants, especially guanidine thiocyanate, primarily present in the RNA lysis buffers. In case of retina derived RNA, both mice strains provided comparable results. Similar observations of slightly lower 260/230 ratios were made in BALB/c mice derived retinal RNA.

**Figure 2:**
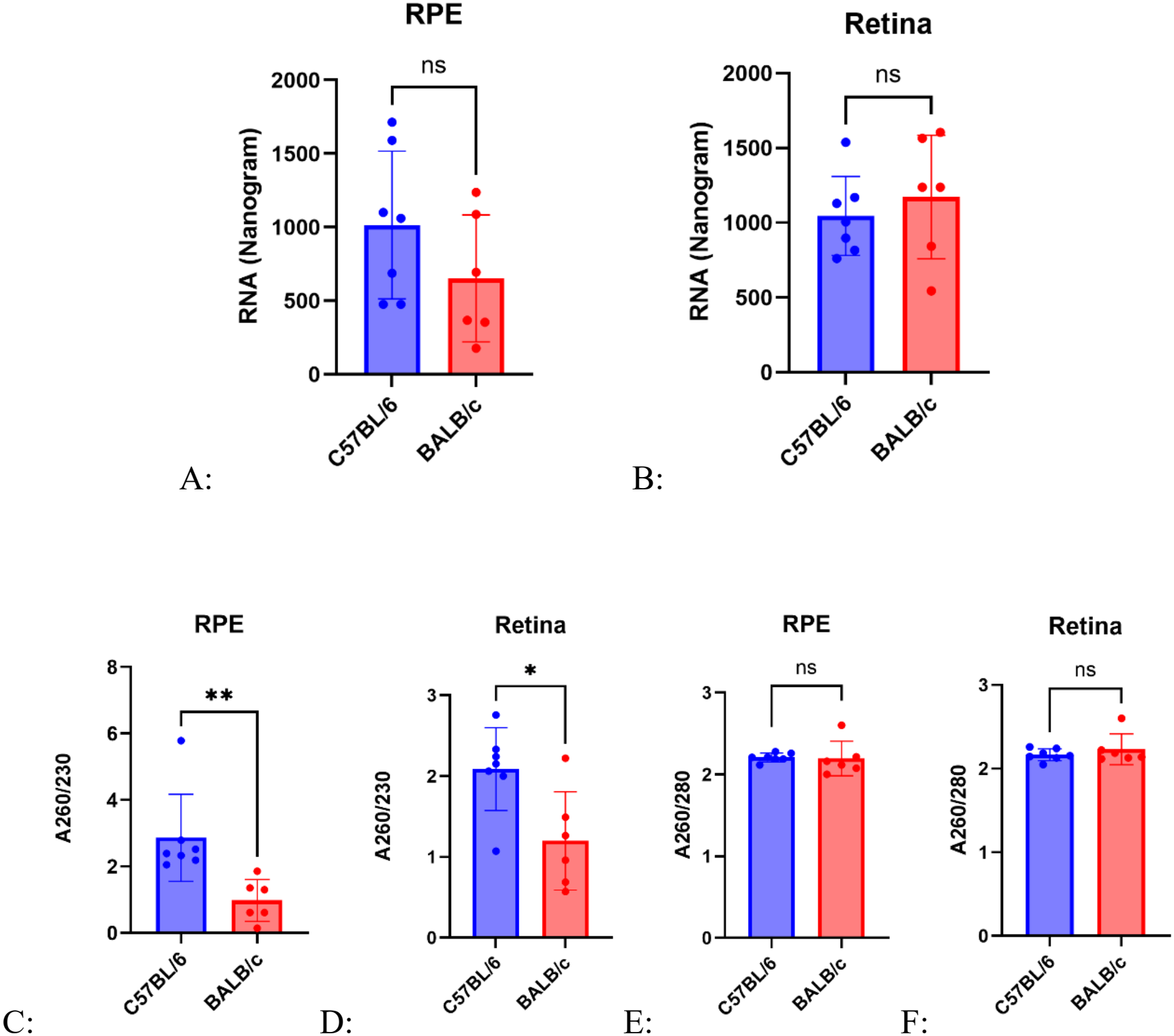
(A-F): Comparison of the yield and purity of obtained RPE and retinal RNA between C57BL/6 and BALB/c. A&B: RPE and retinal derived RNA yield between C57BL/6 and BALB/c, respectively. C&D: A260/230 ratios of RPE and Retinal RNA. E&F: A260/280 ratios of RPE and retinal RNA. Error bars: means ± SD. \*\**P* < 0.01, \**P* < 0.05, Ns: non-significant, Welch’s T test is conducted using GraphPad Prism.

### 3.2. Immunostaining shows clean separation

The immunofluorescent staining with beta-catenin represented the success of our isolation method in RPE removal from the whole choroidal flat-mounts (Figure 3 & 4). The choroidal tissue not used for RNA isolation showed the preserved hexagonal ‘honeycomb’ patterns, corresponding to the unperturbed cellular morphology of RPE cells in the whole flat mounts. Compared to those, the choroid tissue undergoing isolation method showed no such hexagonal patterns. The lack of RPE specific structural integrity observed in the post-isolation choroidal tissue validated the success of our technique in removal of RPE cells. These data were comparable in both BALB/c and C57BL/6 as observed by Figure 3 and 4. In some parts of the tissue, some irregular patches of staining were observed. These signals were most likely occurring due to the exposed choroid capillaries post-isolation.

**Figure 3:**
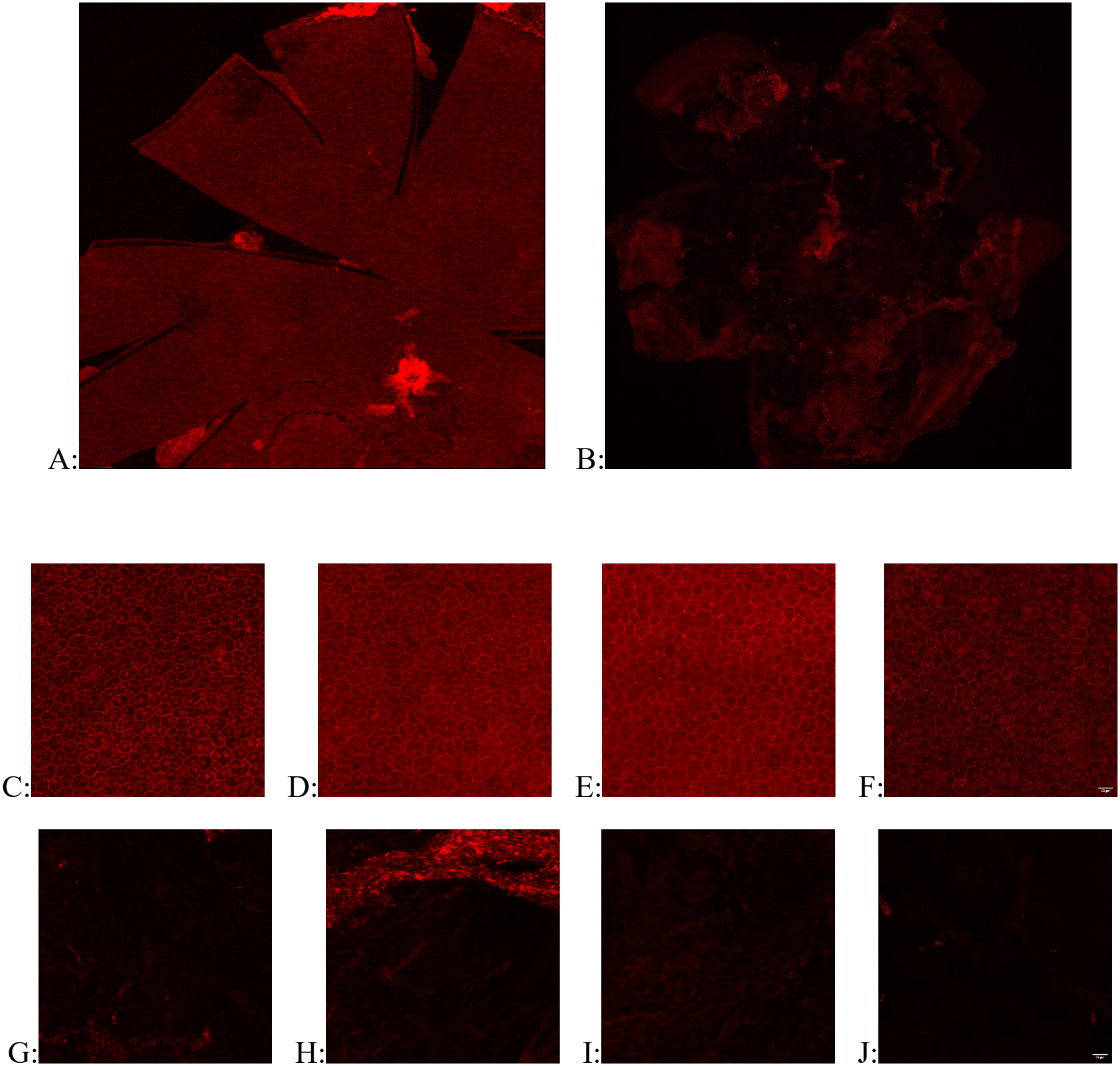
Beta-catenin immunostaining of flat-mount choroid tissue shows RPE removal in C57BL/6. (A-B): Tiled whole flat-mount images of tissues prior (A) and post RNA isolation (B). (C-D): Representative images of intact hexagonal RPE structures in choroidal tissue prior undergoing any isolation, (G-J): Representative images of choroidal tissue post-isolation. Scale bar: 20 um.

**Figure 4:**
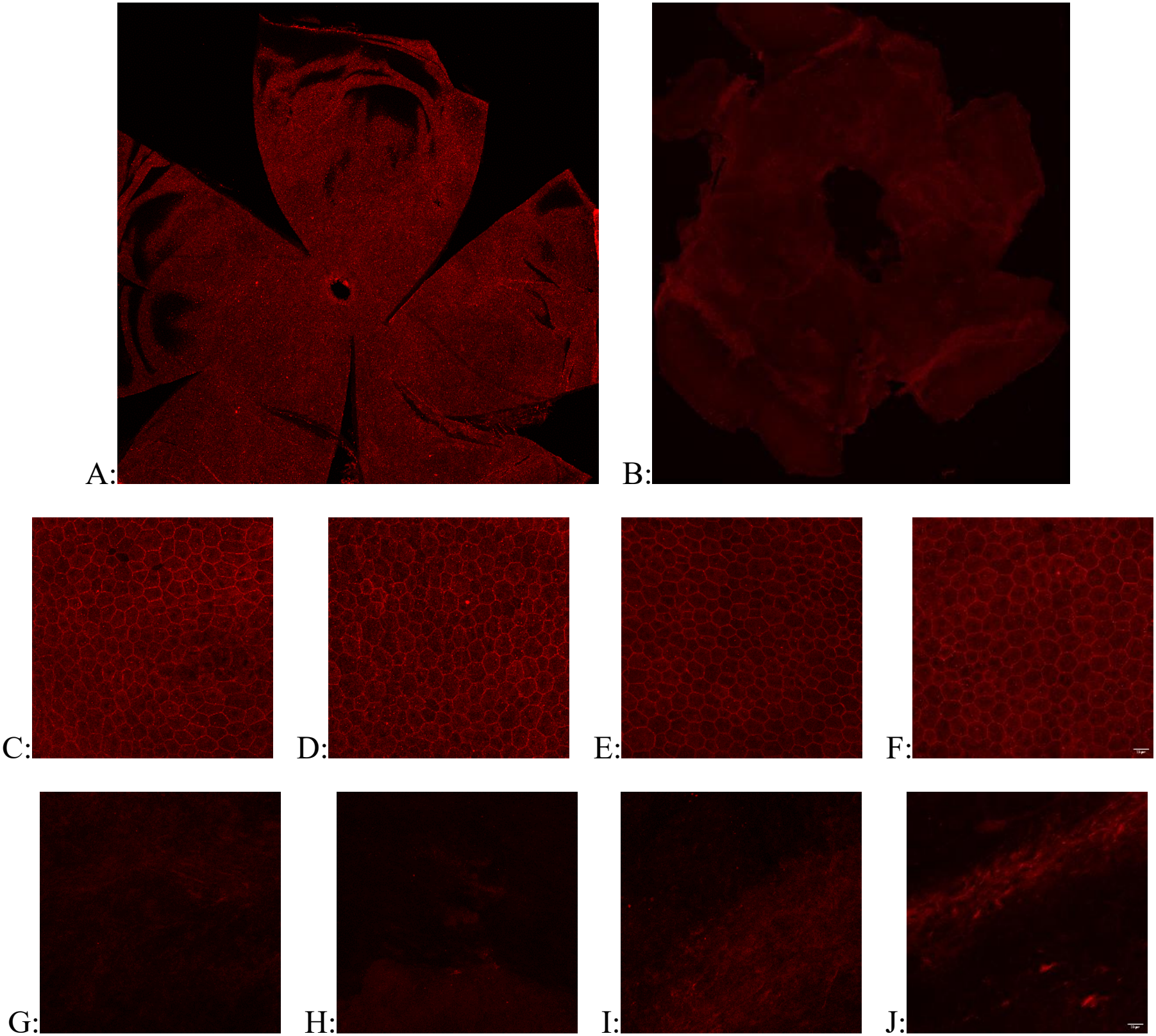
Beta-catenin immunostaining of flat-mount choroid tissue shows RPE removal in BALB/c. (A-B): Tiled whole flat-mount images of tissues prior (A) and post isolation (B). (C-F): Representative images of intact hexagonal RPE structures prior undergoing any isolation. (G-J): Representative images of choroidal tissue post-isolation. Scale bar: 20 um.

### 3.3. Biomarker analysis

The presence of Collagen-6 in the C57BL/6 RPE RNA was minimal compared to the RPE65 (Figure 5). This data validated that the isolated RNA was majorly derived from RPE. But unfortunately, in BALB/c strain, greater contamination of choroid capillaries tissue was observed in the RPE RNA. Ocular pigmentation of murine eye played a role in the dissection process of choroid and retina. The lack of pigmentation of BALB/c choroid and retina made the separation more difficult and impacted the downstream experiments. In both C57BL/6 and BALB/c strains, the retinal RNA samples were found to be predominantly pure as observed by the very low presence of RPE65 compared to Rhodopsin.

**Figure 5:**
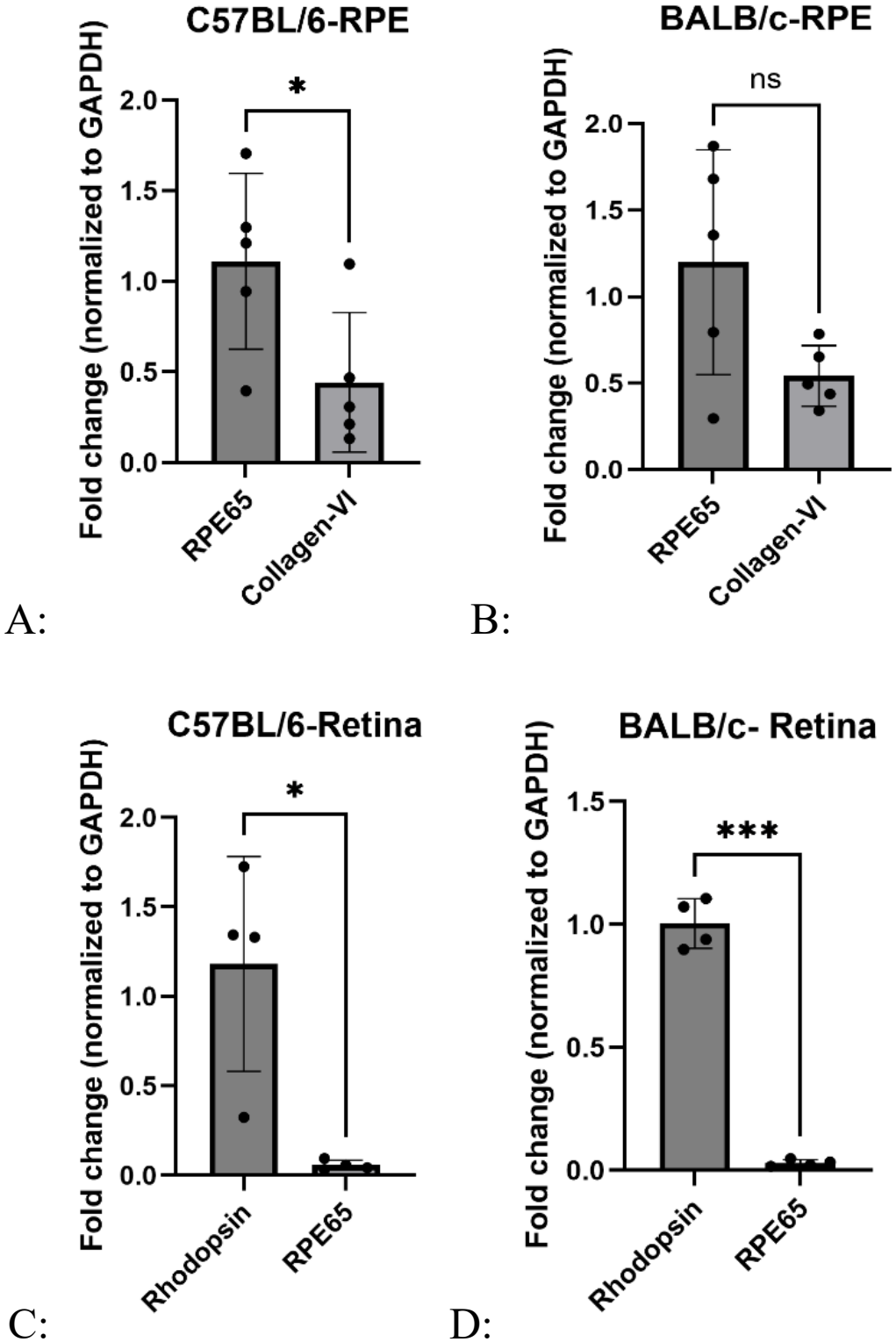
Comparative analysis shows minimal other cellular contamination. A: Significantly different fold change of RPE65 (RPE biomarker) and collagen-VI (choroid capillaries biomarker) in C57BL/6 RPE RNA, B: BALB/c RPE derived RNA shows presence of other cell types. C&D: Minimal RPE-65 present in the retina RNA in C57BL/6 and BALB/c retina, respectively. Error bars: means ± SD. \*\*\**P* < 0.001, \**P* < 0.05, Ns: non-significant, Welch’s T test is conducted using GraphPad Prism.

## 4. Discussion

In this paper we described a rapid method to isolate RNA from both murine RPE and retina simultaneously. This isolation protocol provided a relatively clean separation of RPE and retina across different mice strains. This is the first described complete method of isolating both RPE and Retinal RNA simultaneously. One major advantage of our method is speed. Traditional homogenization methods for RNA isolation often require longer incubation periods on ice [14]. But our method didn’t require longer incubation periods. Our method had some similarities with the standard SIRS method [16]. But unlike the SIRS method, we didn’t pellet the RPE in the RNA-protect reagent. The RNA-protect reagent helped to stabilize the RNA within the tissue. For isolating RNA, we dissected the eye into choroid and retina and homogenized the tissues directly into RLT buffer. This modified method provided us with more flexibility to isolate RNA even after several days post-euthanasia. Different research groups employed modifications to the SIRS method to serve different purposes. For example, a two-step isolation process was implemented to obtain the RPE and sclera-choroidal RNA separately [18]. A combination of SIRS and PRO200 Homogenizer mediated RPE and retinal RNA isolation was also described [19]. That combined method described instant separation of retina and choroidal tissue post euthanasia. Contrary to their technique, we used a smaller hand-held homogenizer for retinal RNA isolation. The use of disposable sterile pestles and ease of using the smaller homogenizer made our method less prone to cross-sample contamination. We also used the same extraction methods for both RPE and retinal RNA isolation; thus, it streamlined our workflow. Based on the qPCR and immunostaining data, our method was found to be a reproducible RNA isolation technique across different mice strains. One limitation of our method was the presence of possible salt contaminants from the lysis buffer as shown by the 260/230 ratios in BALB/c mice RNAs. Mouse eye is considered a low-RNA yielding tissue with less than 10 million cells present per eye [20-21]. The sub-population of RPE consists of approximately 50000 cells per mouse eye [22]. So, the presence of even a miniscule amount of leftover chemical showed up in the 260/230 values of RPE derived RNA. But up to millimolar concentration of this salt was found have no inhibitory effects on PCR as validated by the kit manufacturers. We observed that the lighter pigment of the BALB/c eyes made the process of dissecting the RPE and retina challenging. Darker pigmented C57BL/6 eyes did not have such an issue. Our method can be a useful tool to isolate good quality RNA from small sub populations of ocular cell types. This protocol can help to generate considerable RNA yield to understand disease focused downstream analyses.

## Acknowledgements

The Authors appreciate the funding obtained from LIFT funds to conduct this study. The authors would like to acknowledge Dr. Patrick Kerstein and Dr. Michael Dorrell for their valuable insights on the isolation protocol.

## Author Contribution

SS developed the isolation method. SS designed the experiments and wrote the manuscript. MS supervised the experimental workflow and edited and revised the manuscripts.

